# Root rot by *Phytophthora cinnamomi* shifts the composition and structure of avocado rhizosphere fungal communities

**DOI:** 10.64898/2026.07.10.737851

**Authors:** Rosaura G. Alfaro-García, Alejandro M. Cisneros-Martínez, Violeta Patiño-Conde, Eria A. Rebollar, José A. Guerrero-Analco, Alfonso Méndez-Bravo, Frédérique Reverchon

**Affiliations:** Red de Diversidad Biológica del Occidente Mexicano, Centro Regional del Bajío, Instituto de Ecología, A. C. Av. Lázaro Cárdenas 253, 61600 Pátzcuaro, Michoacán, México; Laboratorio Nacional de Análisis y Síntesis Ecológica, Escuela Nacional de Estudios Superiores, Unidad Morelia, Universidad Nacional Autónoma de México. Antigua carretera a Pátzcuaro 8701, Ex Hacienda de San José de la Huerta, 58190 Morelia, Michoacán, México; Centro de Ciencias Genómicas, Universidad Nacional Autónoma de México. Av. Universidad s/n, 62220 Cuernavaca, Morelos, México; Red de Estudios Moleculares Avanzados, Instituto de Ecología, A.C. Carretera Antigua a Coatepec 351, 91073 Xalapa, Veracruz, México; SECIHTI, Av. Insurgentes Sur 1582, 03940 Ciudad de México, México

**Author notes:** Co-corresponding authors: Frédérique Reverchon, Alfonso Méndez-Bravo.

**Keywords:** Microbial interactions, Opportunistic pathogens, Pathobiome, Saprotroph, Soil-borne diseases

## Abstract

Rhizosphere microbial communities contribute to the growth and health of their host but may be altered by the incidence of soil-borne pathogens. In avocado, the oomycete *Phytophthora cinnamomi*, causal agent of *Phytophthora* root rot (PRR), has been shown to alter rhizosphere bacterial communities, although its effect on fungal communities has seldom been explored. Our objective was thus to determine whether *P. cinnamomi* induced shifts in diversity, composition and co-occurrence networks of fungal communities in the rhizosphere of avocado trees, and to identify potential antagonists of *P. cinnamomi* that could be further considered for disease management. Fungal communities associated with the rhizosphere of asymptomatic and PRR-symptomatic avocado trees were studied through ITS metabarcoding. Although α-diversity metrics were not significantly different between asymptomatic and PRR-symptomatic trees, differences in β-diversity of rhizosphere fungal communities were detected. Moreover, PRR led to the enrichment of saprotrophic taxa and opportunistic pathogens such as *Fusarium*, *Cladosporium* or *Plectosphaerella* in the avocado rhizosphere, which were possibly attracted by the release of resources from necrosed roots. Co-occurrence network analysis revealed that fungal networks in the rhizosphere of PRR-symptomatic trees were more complex and connected than those from asymptomatic trees, suggesting a response of fungal communities to the disturbance caused by the pathogen. Some connector taxa from the PRR-symptomatic networks (*Gibellulopsis*, *Cladorrhinum* or *Mycenella*) were also identified as members of the *P. cinnamomi* pathobiome. Their negative correlations with the pathogen indicate they may act as potential antagonists, which calls for further isolation efforts to confirm their biocontrol activity of PRR.

## 1. Introduction

Rhizosphere microorganisms contribute to their host‘s growth and fitness by improving the uptake of essential nutrients, by stimulating plant immunity or by providing protection against pathogens and abiotic stressors (Berendsen et al., 2012; Park et al., 2023). The rhizosphere is also a suitable habitat for potential plant pathogens, where pathogenic microorganisms can generate a disruption of the eubiosis, defined as the optimal homeostatic state of the microbiota necessary to maintain the overall health of the host, and induce dysbiosis (Paasch and He, 2021). Numerous studies have shown that the incidence of plant pathogens in the rhizosphere can induce shifts in microbial communities, affecting not only their diversity (Mendes et al., 2023), but also their composition (Wang et al., 2022b; Liu et al., 2023) and functions (Batista et al., 2024; Alfaro-García et al., 2026b; Basu et al., 2026). However, these pathogen-induced shifts vary depending on the pathosystem and environmental conditions, thus warranting a deeper understanding of the effects of soil-borne pathogens on rhizosphere microbial communities associated with crops of high economic relevance (Alfaro-García et al., 2026a).

Avocado (*Persea americana* Mill.) is one of the four most traded tropical fruits in the world (FAO, 2026). In 2019, Mexican avocado plantations covered about 234,570 ha and supported a production of 2.3 million tons, of which 1.2 million tons were exported, thus making Mexico the main exporter of this fruit (FAO, 2026; Díaz-Castellanos, 2021). Nevertheless, avocado production faces several challenges such as soil degradation, increased drought due to deforestation and climate change, and the incidence of pests and pathogens (Sánchez-Hernández et al., 2025). One of the most important pathogens affecting avocado orchards globally is *Phytophthora cinnamomi* Rands., the causal agent of Phytophthora root rot (PRR), a disease causing the decay of feeder roots and subsequent tree dieback, often leading to estimated losses orchard losses from 50 to 90 % (Fernández-Pavia et al., 2013). Several works have examined the shifts in microbial communities induced by *P. cinnamomi* in the avocado rhizosphere, with a large focus on bacteria (Yang et al., 2001; Shu et al., 2019; Solís-García et al., 2021; Farooq et al., 2022; Alfaro-García et al., 2026b). However, parallel studies from our group suggests that the effect of *P. cinnamomi* may be more pronounced on rhizosphere fungal communities than on bacterial ones (Reverchon et al., 2023; Alfaro-García et al., 2026b) and that the observed disease symptoms may be induced by the interactions of various pathogenic microorganisms rather than by the sole action of *P. cinnamomi* (Solís-García et al., 2021). In this context, deciphering the interactions between *P. cinnamomi* and fungi in the avocado rhizosphere may represent an important step for disease management.

Such interactions between the main disease causal agent and the rhizosphere microbiota have been studied in other pathosystems to elucidate the “pathobiome”, i.e., the set of microorganisms associated with the plant diseased state (Bass et al., 2019; Alfaro-García et al., 2026a). For example, Guo et al. (2021), characterized the structure and co-occurrence networks of rhizosphere microbial communities of peanut (*Arachis hypogaea* L.) infected by the fungus *Athelia rolfsii*. These authors highlighted that, in infected soils, the relative abundances of *Solicoccozyma*, *Fusarium* and *Humicola* were negatively correlated with *A. rolfsii* abundance, suggesting that these genera could directly or indirectly affect the population dynamics of the pathogen. Dastogeer et al. (2022) identified the pathobiome of *Magnaporthe oryzae* in rice plants and showed that the relative abundance of *M. oryzae* was negatively correlated with that of *Penicillium* and *Desulfobacca* in unhealthy soils, suggesting that these microorganisms could regulate pathogen populations. In cotton, Qiu et al. (2022) elucidated the pathobiome of *Fusarium oxysporum* and found that 32 fungal operational taxonomic units (OTUs) were significantly correlated with the pathogen. In particular, taxa such as *Trichoderma* sp., *Acremonium* sp. or *Fusarium* spp. were negatively associated *F. oxysporum*, suggesting that these microorganisms may provide protection against the pathogen. Such co-occurrence networks may therefore constitute a promising approach to elucidate disease pathobiomes and to identify putative biocontrol agents.

Co-occurrence networks are also promising tools to study potential interactions within the microbiome. Connected microbial communities may be more stable, self-organized, and thus more resilient to perturbations than communities with a lesser degree of connectivity (Dundore-Arias et al., 2023). In the context of soil-borne pathogens, increased connectedness suggests more cooperation within the microbial community (Cardoni et al., 2023), which may mitigate the increase in pathogen populations whilst contributing to suppressing plant infection (Batista et al., 2024). In addition, identifying key taxa in the networks may direct our search for potential biocontrol agents and future experiments aimed at validating their bioactivity *in planta* (Barberán et al., 2012; Gao et al., 2022; Dundore-Arias et al., 2023).

The objectives of this work were thus to assess the impact of PRR on the structure, composition and interactions of rhizosphere fungal communities in avocado, under field conditions. Moreover, we aimed to identify the pathobiome associated with *P. cinnamomi* by using co-occurrence networks, to gain a better understanding of fungal taxa that may be contributing to, or counteracting, the disease.

## 2. Materials and methods

### 2.1. Study site and rhizosphere soil sampling

On April 2023, we selected an orchard located in Peribán, Michoacán (19° 31’16“ N, 102° 24’54” W) under organic management and with records of *P. cinnamomi* incidence in recent years. Soil chemical characteristics of the orchard are presented in Alfaro-García et al. (2026b). Within the orchard, we randomly selected 20 avocado trees, aged between eight and 15 years old. From these trees, 10 were asymptomatic and 10 showed symptoms of PRR, namely defoliation, dead or chlorotic leaves, and small, brittle and/or necrotic roots. The presence of *P. cinnamomi* in the rhizosphere of symptomatic trees was confirmed by subsequent culturing in laboratory conditions (Alfaro-García et al., 2026b). Rhizosphere soil (soil adhered to fine roots) was collected at four points around the trunk at a maximum distance of 50 cm and a 10 cm depth. The four samples obtained per tree were then mixed to obtain one composite sample per tree. All samples were stored at-80 °C until DNA extraction.

### 2.2. DNA extraction and ITS2 rRNA library construction

The DNeasy® PowerSoil® Kit (Qiagen) was used to extract DNA from the rhizosphere soil, following the instructions provided by the manufacturer. The purity and concentration of DNA extracts were verified with an Eppendorf BioSpectrometer®. The construction of sequencing libraries followed the workflow plan outlined in 16S Metagenomic Sequencing Library Preparation of Illumina, (2013) with slights modifications as follows. Polymerase chain reaction 1 (PCR1) was used to amplify the Internal Transcribed Spacer 2 (ITS2) gene and to create ∼400 bp amplicons (Tedersoo et al., 2014). PCR1 was performed with an equimolar pool of primers listed in Supplementary Table 1. 50 µl PCR reactions were performed with 1 µl of template DNA (∼ 30-40 ng µl^-1^), 25 µl Multiplex PCR (Qiagen), 2.5 µl of each primer (2 µM) and 19 µl of MilliQ water. PCR cycling parameters were as follows: 95 °C for 15 min, 94 °C for 3 min; 37 cycles at 94 °C for 40 s, 55 °C for 40s, 72 °C for 1 min; and a final extension step at 72°C for 7 min. Each PCR1 product was purified with ProNex® Size Selective Purification System. Once purified, each ITS rDNA amplicon was assigned a dual index and an Illumina sequencing adapter using the Nextera XT Index kitTM in a second PCR (PCR2). The PCR2 cycling parameters were as follows: 95 °C for 18 min; 8 cycles at 95 °C for 30 s, 55 °C for 30 s, 72 °C for 30 s; and a final extension step at 72 °C for 5 min. PCR2 products were subsequently purified with ProNex® Size Selective Purification System. Then, a pool was made with all the libraries for shipment, normalizing their concentration to 11 nM and being dissolved in TE buffer (Tris-EDTA). The libraries were sequenced at each end (paired-end) with a read length of 2 × 300 bp using the Illumina MiSeqTM system at CD Genomics.

### 2.3. Bioinformatic analysis for ITS2 rDNA sequences

The quality control of all ITS2 amplicon sequences was performed using the FastQC software v.0.12.1. (Andrews, 2010). The composition and structure of fungal community was analyzed with the DADA2 pipeline (*dada2* package, Callahan et al., 2016) in R v.4.3.3. (R Core Team, 2023). Forward and reverse sequences were trimmed to a minimum length of 240 bp (filterAndTrim:truncLen). The maximum number of undetermined bases in a sequence was set to zero (filterAndTrim:maxN) and reads having more than two erroneous base calls were removed (filterAndTrim:maxEE). Afterwards, the sequences were de-replicated. In this step, identical reads were assigned to a single representative read to reduce redundancy and make the following steps more efficient. Then, an error model was generated by learning the specific error-signature for each forward and reverse sequence. After quality processing, inferences were made into Amplicon Sequence Variants (ASVs). Before taxonomic assignment, paired-end sequences were merged (mergePairs) and chimeras were removed (removeBimeraDenovo). The fungal taxonomic assignment was carried out by comparison with the UNITE database (Abarenkov et al., 2024). Singletons, doubletons, mitochondrion, chloroplast and eukaryotic sequences were removed. Data were deposited in the Sequence Read Archive of the NCBI under BioProject PRJNA1363127.

### 2.4. Statistical analysis of rhizosphere fungal community

Before the alpha and beta diversity analysis, the ASVs table was normalized according to rarefaction at the minimum sample sequencing depth using the rarefy_even_depth function implemented in the *Phyloseq* package v.1.46.0 (McMurdie and Holmes, 2013) to minimize the influence of sequencing depth on the results. The α-diversity metrics (observed ASVs, Shannon and Simpson indices) were calculated using the estimate_richness function implemented in *Phyloseq* for asymptomatic and symptomatic avocado trees. The differences between conditions (asymptomatic *vs*. PRR-symptomatic trees) were tested with the Mann-Whitney-Wilcoxon test. Venn diagrams were constructed to find the exclusive and shared ASV in each condition and were plotted using the *ggven* v.0.1.19 package in R (Gao et al., 2021). A Bray-Curtis dissimilarity matrix was calculated and used to build a Non-Metric Multidimensional Scaling (NMDS) using the ordinate and plot_ordination functions of *Phyloseq* package respectively. For statistical comparison of the rhizosphere fungal community between conditions, a permutational multivariate analysis of variance (PERMANOVA) was implemented, using the *vegan* v.2.7.2 package with 999 permutations (Dixon, 2003). To distinguish true biological differences from potential dispersion effects within the samples in both conditions, a permutational analysis of multivariate dispersions (BETADISP) was used with the betadisper and permutest functions with 999 permutations in R. The differential abundance of fungal ASVs between tree conditions was tested with the *DeSeq2* package v.1.42.1 (Love et al., 2014) and statistical comparison was supported by the Wald test. All statistical analyses were performed in the software R and considered significant at *P* < 0.05.

### 2.5. Asymptomatic and PRR-symptomatic co-occurrence network construction

Co-occurrences networks were separately constructed for each condition (asymptomatic and PRR-symptomatic) using the fungal community data. An input matrix (rarefied) was prepared by selecting only the ASVs that met the prevalence (≥ 0.4 %) and abundance (≥ 11.8 %) criteria, ensuring that interactions represented in the networks are most likely to be biologically real and meaningful. Then, to identify the positive/negative correlations between the fungal ASVs, the Sparse Correlations for Compositional data (SparCC) method and the SParse InversE Covariance Estimation for Ecological Association Inference (SPIEC-EASI) method were implemented in the R package *SpiecEasi* v1.1.3. (Kurtz et al., 2015), for both tree conditions. Random networks were also built for each tree condition to test the significance of the empirical networks. Properties of empirical and random networks were calculated using the R package *igraph* v.2.2.2 (Kolaczyk and Csárdi, 2014) and the means and standard deviation were calculated for statistical comparisons (Wilcoxon test). Significant correlations (r ≥ 0.5, *P* < 0.05) were used to visualize the networks using Cytoscape v.3.10.4. (Shannon et al., 2003).

Keystone fungal taxa were defined based on their Zi-score (within-module connectivity, measures how well a node is connected to other nodes in its own module) and Pi-score (among-module connectivity, measures how well a node is connected to nodes in different modules) (Guimera and Amaral 2005; Deng et al., 2012). Both Zi and Pi scores were calculated using the *igraph* package in R and used to define the nodes as “Network hub” (highly connected nodes within the entire network; Zi > 2.5, Pi > 0.62), “Module hub” (highly connected nodes within modules; Zi > 2.5, Pi ≤ 0.62), “Connectors” (nodes connecting modules; Zi ≤ 2.5, Pi ≥ 0.62) and “Peripherals” (nodes connecting modules with few external connections; Zi < 2.5, Pi < 0.62). Only fungal taxa fitting the first three categories were considered as potential keystone taxa.

### 2.6. Co-occurrence network to identify the Phytophthora cinnamomi pathobiome

The members of the *P. cinnamomi* pathobiome were identified by constructing a co-occurrence network based on taxa abundance. First, a taxonomic assignment of *P. cinnamomi* was performed, using the “sh_general_release_dynamic_s_all_04.04.2024.fasta” from UNITE database. Then, co-occurrence networks were constructed with the SparCC method (Friedman and Alm, 2012), utilizing the SCNIC (Sparse Co-occurrence Network Investigation for Compositional data) algorithm, a tool available in Qiime2 (Shaffer et al., 2023). For visualizing purposes, only positive and negative correlations with statistical support *P* < 0.05 and a correlation with r ≥ 0.5 were selected; all fungal genera that met these correlation parameters were considered as forming the *P. cinnamomi* pathobiome.

## 3. Results

### 3.1. PRR affects the structure but not the α-diversity of rhizosphere fungal communities in avocado

A total of 879,228 high quality reads were obtained after quality filtering and corresponded to 990 ASVs. Rarefaction curves showed that rhizosphere soil samples of both asymptomatic and PRR-symptomatic avocado trees reached a plateau, indicating that most fungal ASVs were detected in the sampling (Supplementary Figure 1).

Fungal richness and α-diversity indices did not differ significantly in the rhizosphere of asymptomatic and PRR-symptomatic avocado trees (Supplementary Figure 2). However, fungal community structure (β-diversity) was significantly different in the rhizosphere of asymptomatic and PRR-symptomatic avocado trees (PERMANOVA, df = 1, F= 1.87, *P* = 0.026) (Figure 1A). Moreover, the NMDS showed that the fungal community associated with PRR-symptomatic trees was more dispersed than asymptomatic trees. The permutational analysis of multivariate dispersions (BETADISP, *P* = 0.018) confirmed that fungal rhizosphere communities of PRR-symptomatic trees exhibited greater heterogeneity than those associated with asymptomatic trees in their species composition intra-groups.

**Figure 1.**
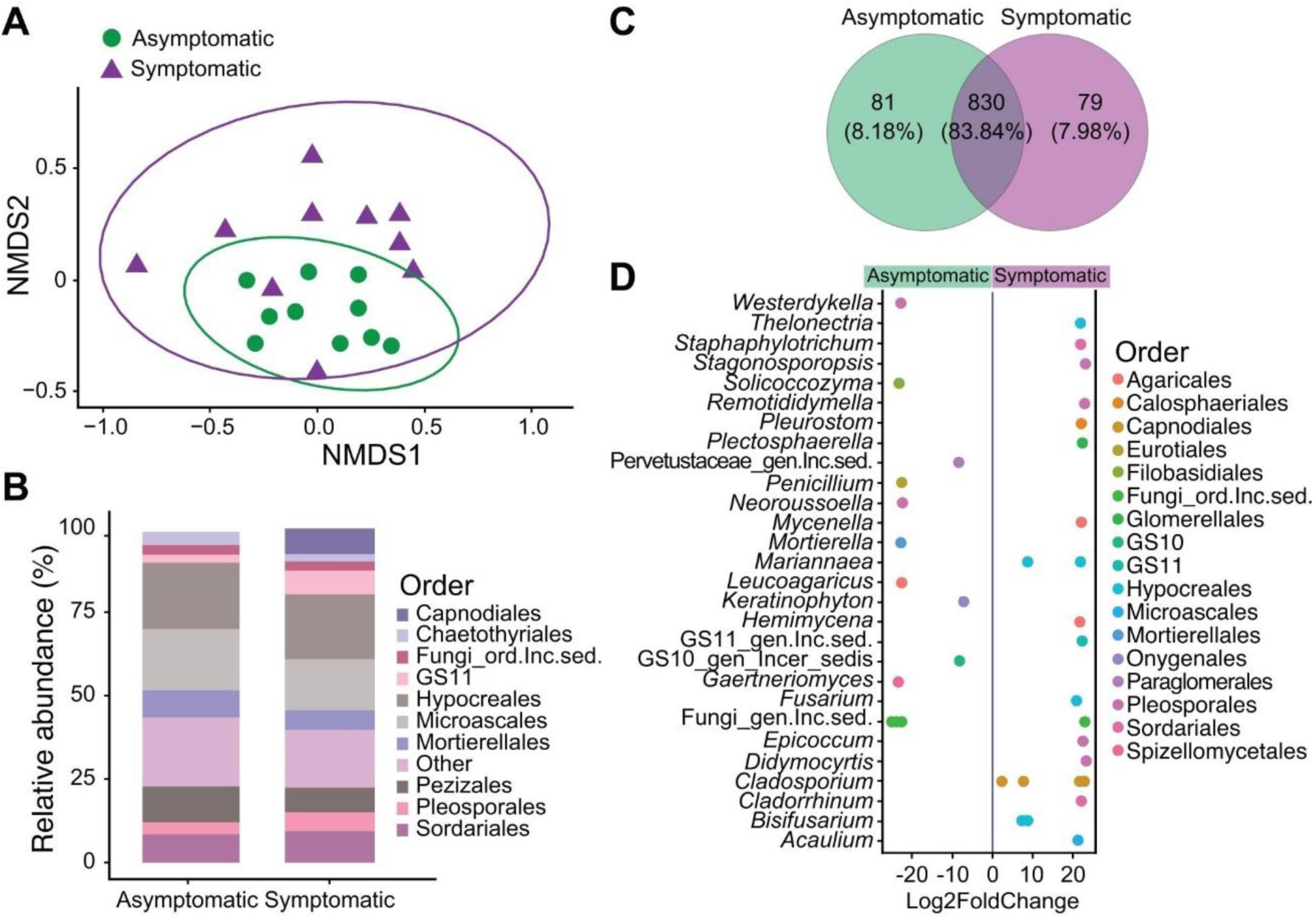
A) Non-metric multidimensional scaling (NMDS) plot based on Bray-Curtis distance of the rhizosphere fungal community structure in asymptomatic and PRR-symptomatic avocado trees (stress value = 0.15, n = 10). B) Relative abundance of the principal fungal orders in each tree condition. The category “others” represents orders with a relative abundance < 1 %. C) Venn diagram of shared and exclusive fungal ASVs in the rhizosphere of asymptomatic and PRR-symptomatic avocado trees (n = 10). D) Differential abundance of fungal ASVs assigned to genera associated with asymptomatic (negative values) and PRR-symptomatic (positive values) avocado trees. According to the Wald test, values of Log2FoldChange ≠ 0 and *P* < 0.05 are considered as differentially abundant.

The rhizosphere of asymptomatic avocado trees was dominated by the fungal orders Hypocreales (19.75 %), Microascales (18.32 %), Pezizales (10.76 %), Sordariales (8.47 %) and Agaricales (2.33 %) orders, while in the rhizosphere of PRR-symptomatic avocado trees, Hypocreales (19.36 %), Microascales (15.36 %), Sordariales (9.36 %), Pezizales (7.39 %) and Capnodiales (7.68 %) were the most abundant orders (Figure 1B).

The asymptomatic and PRR-symptomatic trees shared 84.4 % of fungal ASVs in their rhizosphere (Figure 1C). Each tree condition had the same percentage of unique fungal ASVs (7.8 %). Analysis of differential abundance of rhizosphere fungi showed that ASVs assigned to the genera Fungi_gen_Incertae_sedis (Log2FoldChange =-24.91), *Gaertneriomyces* (Log2FoldChange =-23.57), *Solicoccozyma* (Log2FoldChange =-23.39), *Mortierella* (Log2FoldChange =-22.95) and *Penicillium* (Log2FoldChange =-22.91) were differentially enriched in the rhizosphere of asymptomatic avocado trees. Contrastingly, the fungal ASVs assigned to *Cladosporium* (Log2FoldChange = 22.59), *Didymocyrtis* (Log2FoldChange = 23.11), *Stagonosporopsis* (Log2FoldChange = 23.08), *Remotididymella* (Log2FoldChange = 22.98) and *Fusarium* (Log2FoldChange = 20.00) (Figure 1D) were enriched in the rhizosphere of PRR-symptomatic avocado trees.

### 3.2. Asymptomatic and PRR-symptomatic fungal co-occurrence networks

Co-occurrence fungal networks analysis revealed that fungal network in the rhizosphere of asymptomatic avocado trees consisted of 100 nodes and 435 edges (Figure 2A) whilst that of PRR-symptomatic trees consisted of 104 nodes and 559 edges (Figure 2B). Moreover, fungal network in the rhizosphere of PRR-symptomatic trees represented a larger number of connections (edges) between nodes and an increased density of connections, displaying a more complex structure than the network in the rhizosphere of asymptomatic trees (Table 1). The fungal network of asymptomatic trees exhibited a slightly higher proportion of positive correlations than that of PRR-symptomatic trees (51.03 % vs. 48.83 %).

**Figure 2.**
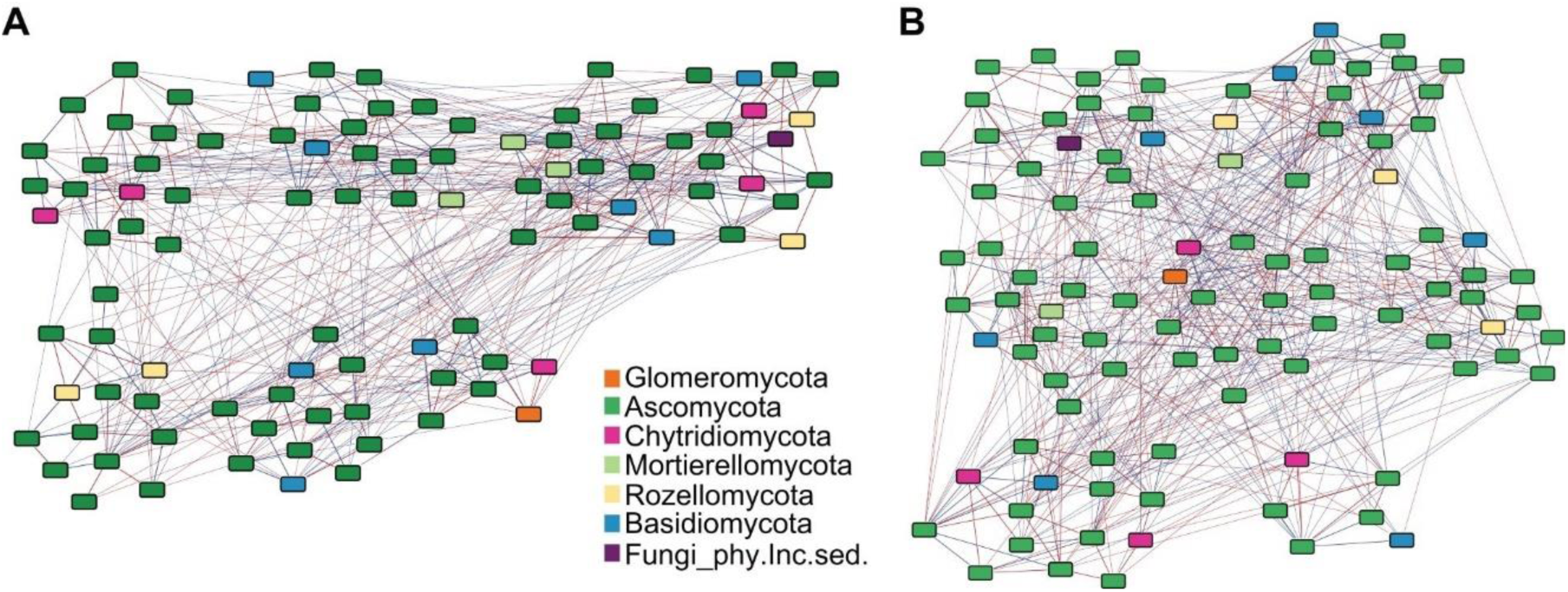
Co-occurrence networks of rhizosphere fungal communities associated with A) asymptomatic and B) PRR-symptomatic avocado trees. The nodes represent an ASV in datasets for each tree condition. Colors represent different fungal phyla. Edges represent significant correlations (r ≥ 0.5 and *P* < 0.05) between nodes. Blue edges show positive correlations and red edges show negative correlations.

**Table 1.** Network properties of asymptomatic and PRR-symptomatic fungal rhizosphere communities.

| Network properties | Empirical networks |  | Wilcoxon test | Random networks |  |
| --- | --- | --- | --- | --- | --- |
|  | Asymptomatic | PRR-symptomatic |  | Asymptomatic | PRR-symptomatic |
| Degree | 8.70 | 10.75 | **W= 3558.5,**<br>**P < 0.001** | 5.26 | 7.02 |
| Clustering coefficient | 0.0927 | 0.1295 | **W= 3317,**<br>**P < 0.001** | 0.0513 | 0.0688 |
| Betweenness | 0.0136 | 0.0116 | W= 5851,<br>P = 0.1228 | 0.0196 | 0.0154 |
| Average shortest path | 2.31 | 2.17 | **W= 7693,**<br>**P < 0.001** | 2.89 | 2.53 |
| Edge density | 0.0878 | 0.1043 |  | 0.0531 | 0.078 |
| Modularity | 0.2770 | 0.2593 |  | 0.3535 | 0.3163 |

The network structures in both tree conditions were verified as reliable and non-random by comparing them with random co-occurrence networks (Table 1). Compared to the random networks, both empirical networks display greater connectivity, clustering and compactness, but less modularity. In the asymptomatic trees, the network of the rhizosphere fungal community showed a lower clustering coefficient than that of the PRR-symptomatic trees, suggesting that microbial interactions are stronger in the symptomatic condition. Also, the values of the average shortest path in the fungal network from PRR-symptomatic trees were lower than those in the asymptomatic trees, indicating potential greater robustness of microbial co-occurrence networks.

Neither network hubs nor module hubs were detected in the fungal network from asymptomatic trees (Figure 3A). However, a network hub (0.96 %, Figure 3B) belonging to the *Botryotrichum* genus was identified in the network from PRR-symptomatic trees. Eighty-seven nodes were defined as connectors in the asymptomatic network, of which 22 were formed by taxa exclusive of this condition (e.g., *Agaricus*, *Aspergillus*, *Colletotrichum*, *Leucoagaricus* and *Westerdykella*). Conversely, 85 nodes were classified as connectors in the PRR-symptomatic network, of which 22 were exclusive of this condition. Node connectors of the PRR-symptomatic network include the fungal taxa *Metarhizium*, *Gibellulopsis*, *Epicoccum*, *Cladorrhinum*, *Didymocyrtis*, *Mycenella* and *Nigrospora* (Supplementary Table 2). Consistently, connector nodes had a proportionally larger representation in the asymptomatic network (Figure 3), indicating a greater connectivity among modules in the asymptomatic condition.

**Figure 3.**
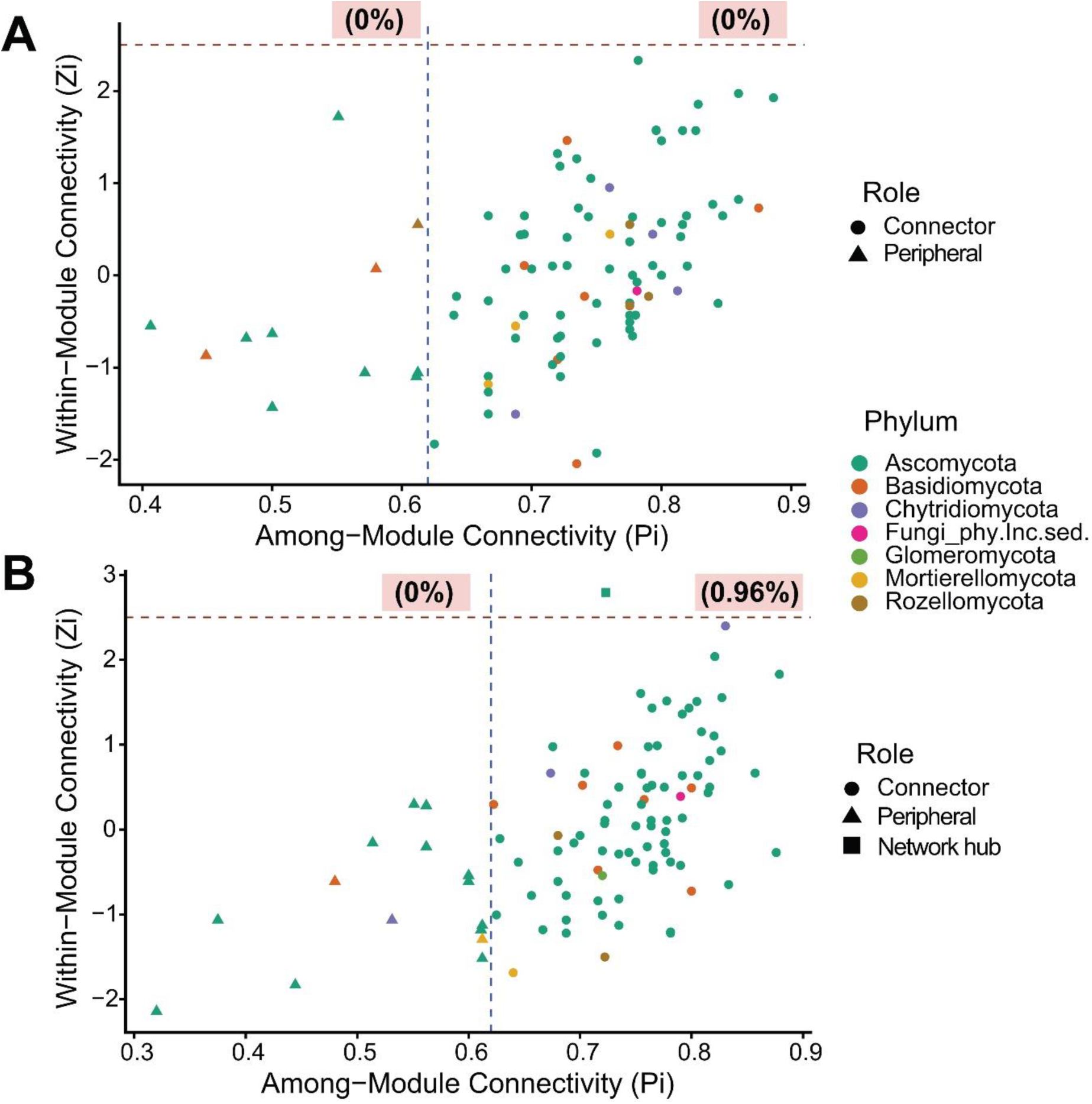
Zi-Pi plots based on ASVs properties and roles in the co-occurrence networks from A) asymptomatic and B) PRR-symptomatic avocado trees. Roles of fungal nodes are shown at the phylum level in the networks. The threshold values for Zi and Pi for categorizing the ASVs roles were 2.5 (red dashed line) and 0.62 (blue dashed line), respectively.

### 3.3. Identification of the pathobiome of Phytophthora cinnamomi

The co-occurrence network allowed the identification of 25 fungal genera as members of the pathobiome of *P. cinnamomi* in the rhizosphere of PRR-symptomatic avocado trees. From these, 12 fungal genera were positively correlated and 13 were negatively correlated with the pathogen (Figure 4). The fungal ASVs assigned to *Leucocoprinus*, *Hannaella* and GS10_gen_Incertae_sedis showed the strongest positive correlations (r = 0.57, 0.57 and 0.55 respectively) with the pathogen, while *Neopyrenochaeta*, *Cladorrhinum* and *Idriella* showed the strongest negative correlations (r =-0.77,-0.58 and-0.52 respectively) with *P. cinnamomi* within the pathobiome (Figure 4).

**Figure 4.**
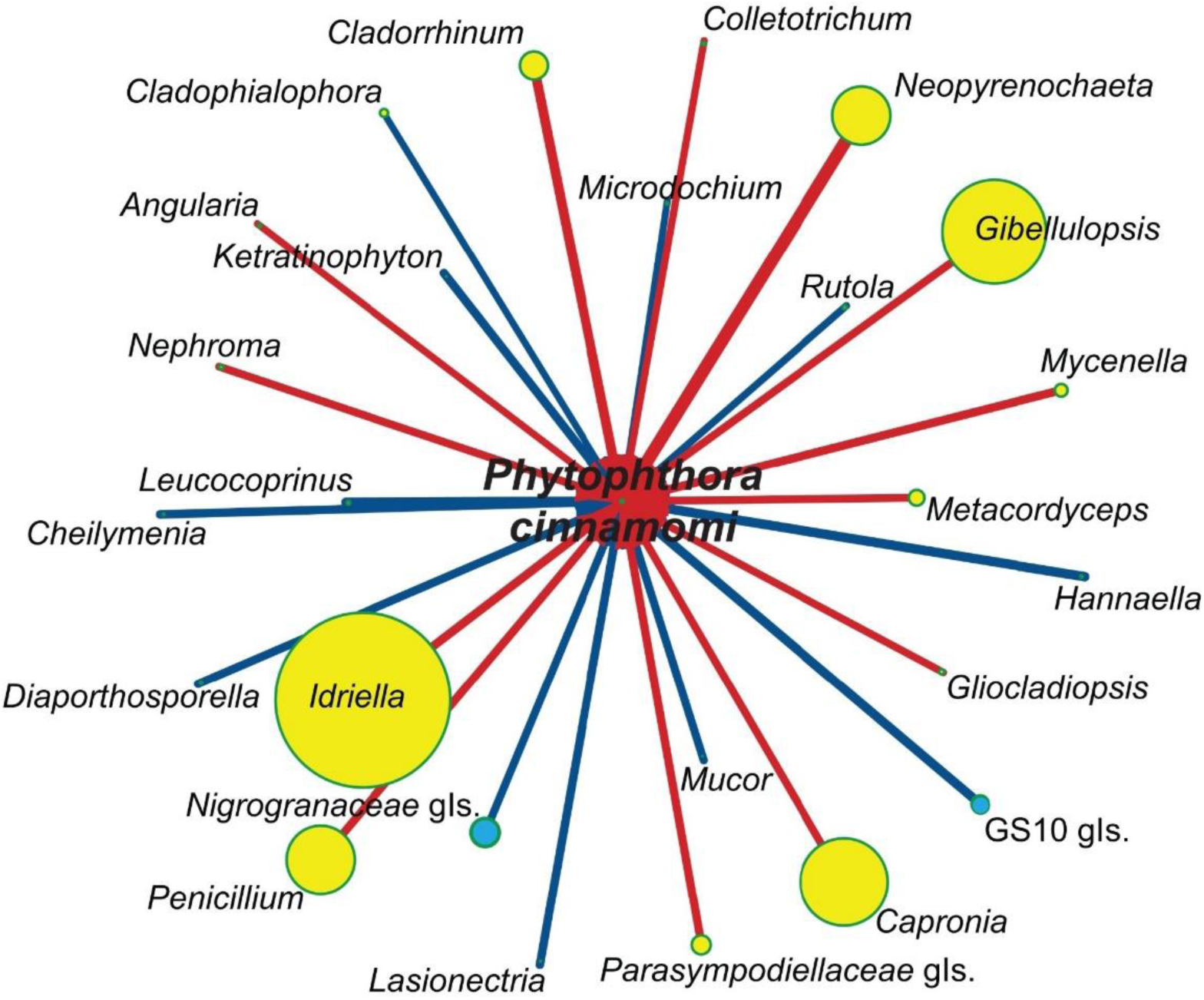
Pathobiome of *Phytophthora cinnamomi*. The node size is related to the relative abundances of the fungal genera. Blue vertices indicate positive correlations and red vertices indicate negative correlations of the fungi with the pathogen. The thickness of every vertex corresponds to the correlation value (r > 0.5). All correlations presented in the pathobiome were significant (*P* < 0.05) based on a correlation matrix constructed with the SparCC method (n = 10).

## 4. Discussion

The objective of our study was to characterize the rhizosphere fungal communities associated with asymptomatic and PRR-symptomatic avocado trees to determine whether the presence of *P. cinnamomi* could be associated with changes in fungal composition and structure. Our results indicated that, although the oomycete did not affect the α-diversity metrics of fungal communities in the avocado rhizosphere, it altered their structure (β-diversity) and induced shifts in their composition and potential interactions. In particular, the presence of the pathogen induced shifts towards a more dispersed fungal community in the rhizosphere, a pattern already described for other pathosystems such as *Ralstonia solanacearum*-tomato or *Xanthomonas oryzae*-rice (Wei et al., 2018; Jobert et al., 2025), indicating a possible dysbiosis and stress-induced destabilization of rhizosphere fungal communities.

We detected an enrichment of fungal ASVs belonging to pathogenic and saprotrophic genera in the rhizosphere of PRR-symptomatic trees, such as *Cladosporium* and *Fusarium*. *Cladosporium* spp. have been described as opportunistic pathogens of diverse plants (e.g., cereals, Curcubitaceae or Solanaceae, Ogórek et al., 2012). Reports include pathogenicity in avocado fruits (Ramírez-Mendoza et al., 2025), although *Cladosporium* spp. were also detected as endophytes in healthy avocado trees (Shetty et al., 2016; Sánchez-Hernández et al., 2025). Similarly, *Fusarium* spp. are ubiquitous saprotrophs and pathogens that have been reported to cause root rot symptoms in a wide variety of crops and wild species (Burgess and Bryden, 2012). In avocado, *Fusarium* spp. are causal agents of root rot (Olalde-Lira et al., 2020; Wang et al., 2025) and were described by Solís-García et al. (2021) as opportunistic pathogens associated with PRR in orchards of Veracruz, Mexico. The enrichment of fungal ASVs belonging to the *Cladosporium* and *Fusarium* genus genera in the rhizosphere of PRR-symptomatic trees may thus confirm the attraction of opportunistic pathogens by the substances released by necrosed roots (Solís-García et al., 2021). Although not all *Cladosporium* and *Fusarium* species may be phytopathogens, their vast array of enzymes and saprotrophic abilities probably explains their enrichment in the vicinity of rotten roots (Steinberg et al., 2016; Moharram et al., 2022). Furthermore, both *Cladosporium* and *Fusarium* have been described as pathogen antagonists and putative biocontrol agents (Chaibub et al., 2016; Nieves-Campos et al., 2024). As recently recommended by Alfaro-García et al. (2026a), further experiments should thus focus on elucidating whether these fungal taxa contribute to the disease or are merely attracted by the nutrient-rich environment favored by root rot.

Another enriched ASV in the rhizosphere of PRR-symptomatic trees belonged to *Plectosphaerella*, a genus known to include necrotrophic pathogens. *Plectosphaerella plurivora*, *P. cucumerina* and *P. melonis* were isolated from tomato, watermelon, melon, bell pepper, asparagus, parsley, ginseng and cabbages (Carlucci et al., 2012; Li et al., 2017; Han et al., 2020), causing root or collar rot disease in all instances. Interestingly, *Plectosphaerella*, as well as *Fusarium*, was also found to be consistently enriched in the rhizosphere of avocado trees infected by *Dematophora necatrix*, causal agent of white root rot (Magagula et al., 2025). However, neither of these genera appeared in the *P. cinnamomi* pathobiome network constructed in our study, although they were identified as members of the *P. cinnamomi* pathobiome in our *in vitro* infection experiment (Reverchon et al., 2023), suggesting the importance of the infection stage and pathogen inoculum density in regulating the pathobiome.

Other fungal genera with ASVs enriched in the PRR-symptomatic condition are *Pleurostoma*, *Mycenella* and *Epicoccum*. Members of the cluster *Pleurostoma*-*Pleurostomophora* are wood-inhabiting fungi that may act as pathogens (Vijaykrishna et al., 2004). *Pleurostomophora richardsiae* is implicated in the decline and branch dieback in olive trees (Carlucci et al., 2013; Ivic et al., 2018; Lawrence et al., 2021) and was also reported as pathogen of grapevines (Carlucci et al., 2015). The *Mycenella* genus comprises agaricoid fungi with saprophytic lifestyle and was reported as one of the most abundant genera in soil samples with presence of *Hymenoscyphus fraxineus*, causal agent of ash dieback (Lysenko et al., 2024). *Mycenella* was identified here as a member of the *P. cinnamomi* pathobiome, although its abundance was negatively correlated with that of the pathogen. In turn, *Epicoccum* is a ubiquitous genus with saprophytic and endophytic lifestyles that has been reported to produce a wide array of secondary metabolites such as polyketides and diketopiperazines with antimicrobial, anticancer and antioxidant activity (Braga et al., 2018), and has recently been described as an avocado endophyte (Sánchez-Hernández et al., 2025). More importantly, its antagonistic activity against soil-borne oomycetes, and in particular against *Phytophthora* spp., points at a possible recruitment of *Epicoccum* by infected avocado trees (Brown et al., 1987; Hashem and Ali, 2004; Li et al., 2013). However, some species have been described as plant pathogens, as *Epicoccum nigrum* in melon and *E. sorghinum* in tobacco (Yuan et al., 2016), sorghum (Oliveira et al., 2017) and pink woodsorrel (Chen et al., 2017). Isolation efforts should thus be made to retrieve *Epicoccum* strains from the rhizosphere of PRR-symptomatic trees and determine whether they contribute to the defense strategy of avocado trees or act as possible avocado pathogens, through *in planta* experiments.

On the other hand, ASVs from the fungal genera *Penicillium*, *Mortierella* and *Solicoccozyma* were significantly enriched in the rhizosphere of asymptomatic trees. *Penicillium* and *Mortierella* are two genera that have been reported by Nieves-Campos et al. (2024) with strong *in vitro* antagonistic activity against *P. cinnamomi*. *Penicillium* was also identified here as a member of the *P. cinnamomi* pathobiome, which relative abundance correlated negatively with that of the pathogen. Its antagonistic activity against different species of *Phytophthora* has been demonstrated by several studies. For example, *Penicillium funiculosum* acted as a biocontrol agent of *P. cinnamomi* in azalea and of *Phytophthora parasitica* in sweet orange, reducing root rot symptoms and allowing the growth of new shoots (Fang and Tsao, 1995). Ai et al. (2023) found that *Penicillium oxalicum* inhibited the growth of *Phytophthora cactorum* by up to 74.4 % in co-culture. Although *Mortierella* was not identified as a member of the pathobiome here, it was shown in other studies to benefit plant health through its capacity to increase nutrient availability in rhizosphere soils and to reduce the symptoms caused by the soil-borne pathogen *Fusarium oxysporum* in ginseng (Wang et al., 2022a). However, *Mortierella* has also been described as an opportunistic pathogen in avocado trees (Hernández Pérez et al., 2018; Solís-García et al., 2021), which warrants further research of the particular ASV identified here.

Other fungal genera were identified as members of the *P. cinnamomi* pathobiome, correlating negatively with the abundance of the pathogen, although they were not detected by the differential abundance analysis as significantly enriched in the rhizosphere of symptomatic trees. Nevertheless, their potential as putative biocontrol agents should not be discarded. In particular, *Neopyrenochaeta* presented the strongest negative correlation with *P. cinnamomi*. Since this genus was also detected in our parallel greenhouse infection study as part of the pathobiome of *P. cinnamomi* in avocado (Reverchon et al., 2023), subsequent isolation efforts should be made to further investigate the biocontrol potential of *Neopyrenochaeta* strains, as this genus has not yet been tested against phytopathogens, to the best of our knowledge. Potential plant pathogenicity of *Neopyrenochaeta* isolates should also be tested. *Cladorrhinum* and *Idriella*, other fungal taxa negatively correlated with *P. cinnamomi* in the pathobiome network have been reported as antagonists of several plants pathogens. Lewis et al. (1995) showed that *Cladorrhinum foecundissimum* significantly reduced the growth of *Rhizoctonia solani* in cotton plants. Using the same pathosystem, Gasoni and de Gurfinkel (2009) showed that two strains of *C. foecundissimum* reduced the symptom incidence of root rot by up to 63.4 %. Collectively, species of *Cladorrhinum* display great potential for biocontrol and plant growth promotion (Martin et al., 2019). In turn, the inoculation of barley with *Idriella bolleyi* over six years led to a 16 % reduction of the severity of common root rot caused by *Bipolaris sorokiniana* and an average yield increase of 4 % (Duczek, 1997). Liljeroth and Bryngelsson (2002) also found that seed and root inoculation of barley with *I. bolleyi* resulted in reduced root rot symptoms, due to the induction of the plant systemic resistance. Collectively, these findings warrant future work to focus on the biocontrol activity of *Cladorrhinum* and *Idriella* against PRR in avocado.

The presence of *P. cinnamomi* also modulated fungal interactions within the avocado rhizosphere. Co-occurrence network analysis evidenced that nodes in networks from the PRR-symptomatic avocado trees were connected with more edges, displayed greater values of degree, clustering coefficient and a lower average shortest path, thus suggesting that fungal communities in the rhizosphere of PRR-symptomatic avocado trees are more complex than those of asymptomatic trees. The increased complexity of microbial communities in diseased soils and roots has been registered in several pathosystems, such as strawberry plants suffering from root rot (Zhang et al., 2023), olive trees infected by *Verticillium dahliae* (Fernández-Gonzalez et al., 2020) and in rice plants infected by *Xanthomonas oryzae* (Jiang et al., 2023). This increase in network complexity may imply a faster response to environmental disturbances such as pathogen infestation, and an increased stability of the fungal rhizosphere communities in *P. cinnamomi* infected roots (Jiang et al., 2023). Moreover, fungal communities in the rhizosphere of PRR-symptomatic avocado trees were less cooperative as the proportion of negative connections increased when compared with asymptomatic networks. These negative correlations could reflect an increase in competition for nutrients released by the necrosed roots, principally carbon resources, which coincides with the enrichment of saprotrophic fungal genera such as *Epicoccum*, *Mycenella*, *Plectosphaerella* in the rhizosphere of PRR-symptomatic trees. Competition for resources has also been seen described as a signal of microbial community stability, since it indicates the presence of metabolically redundant species within the community, which in turn could restrict microbial pathogen overgrowth (Wang and Kuzyakov, 2024). Consistently, the rhizomicrobiomes of eggplants or chili pepper harboring more negative interactions displayed the greatest resistance to *Ralstonia solanacearum* invasion and to *Fusarium* wilt (Gao et al., 2021; Jiang et al., 2022), respectively.

Among the nodes categorized as connectors in the PRR-symptomatic network and identified as exclusive of the symptomatic condition were *Mycenella*, *Cladorrhinum* and *Gibellulopsis*, which were also identified as members of the *P. cinnamomi* pathobiome, where they correlated negatively with the pathogen, suggesting that these taxa could be potential antagonists of *P. cinnamomi* and key taxa for the stability of the fungal community when faced with pathogen invasion. Another identified connector taxon which was unique to the PRR-symptomatic network was *Metarhizium*, a genus well-known for enhancing plant health and as a biological control agent of insect pests (St. Leger and Wang, 2020; Stone and Bidochka, 2020), which could also be key for maintaining the structure and function of the network in the presence of *P. cinnamomi*.

## 5. Conclusions

Although PRR did not induce changes in α-diversity in fungal communities associated with the avocado rhizosphere, the disease led to changes in community structure and more dispersed fungal communities, favoring the enrichment of opportunistic fungal pathogens and saprotrophic fungi in the rhizosphere of PRR-symptomatic trees. Moreover, co-occurrence network analyses showed that fungal communities in the rhizosphere of PRR-symptomatic trees were more complex than those in asymptomatic trees, which suggests a response to the pathogen presence. We elucidated the pathobiome of *P. cinnamomi* and showed that genera such as *Penicillium*, *Neopyrenochaeta*, *Cladorrhinum* and *Idriella* may act as antagonists of the pathogen in the avocado rhizosphere. Although this study only constitutes a snapshot of the dynamic changes avocado rhizosphere fungal communities may undergo when the host tree is confronted to PRR, our findings highlight the importance of pathobiome studies to narrow our search for promising microbial biocontrol agents of PRR in avocado.

## Acknowledgements

This study was financially supported by project FOSEC-SEP-CONAHCYT (SECIHTI), grant number A1-S-30794 and by the CONAHCYT(SECIHTI)-PRONACES-PPE grant (project “PERSEA”, 322772). We thank SECIHTI for the PhD scholarship granted to R.A.- G. (CVU 932217). We would like to thank Alejandra Mondragón-Flores and Silvia Fernández-Pavía for their help with orchard selection and Ing. Victor for granting us access to the orchard. We are grateful to Daniel Sánchez-Hernández, Alejandra Mondragón-Flores and Alejandro Soto-Plancarte for their help with sampling, to Laura Piñon and Eduardo Thome, for their help with the sequencing libraries construction, and to Diego Isla López for his technical assistance with computational tools at Cluster-LANASE, ENES-Morelia, UNAM.

## Data availability

Data are submitted in the SRA database of the NCBI.

