## Supplementary Figures 1 and 2 for "Root rot by *Phytophthora cinnamomi* shifts the composition and structure of avocado rhizosphere fungal communities"

**Supplementary Figure 1. Rarefaction curves of fungal ASVs in the rhizosphere of asymptomatic and PRR-symptomatic avocado trees (n = 10).**

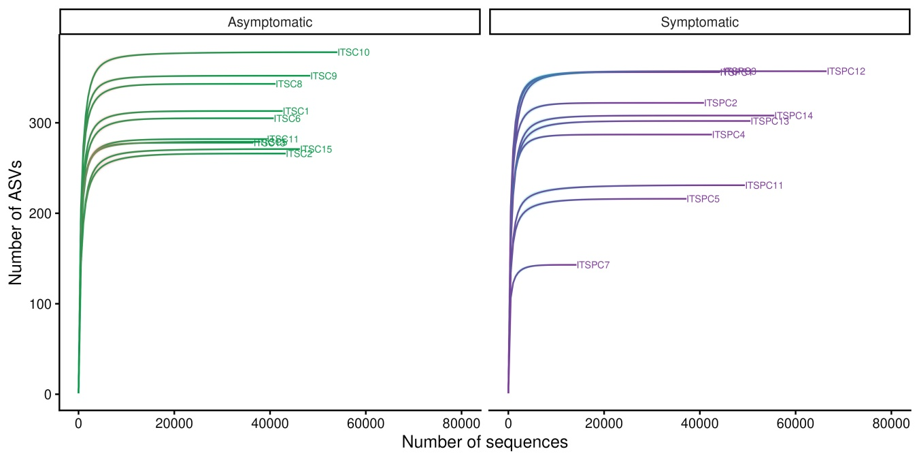

**Supplementary Figure 2. Alpha diversity metrics of rhizosphere fungal communities associated with asymptomatic and PRR-symptomatic avocado trees. W and P value were calculated with the Mann-Whitney-Wilcoxon test (n = 10).**

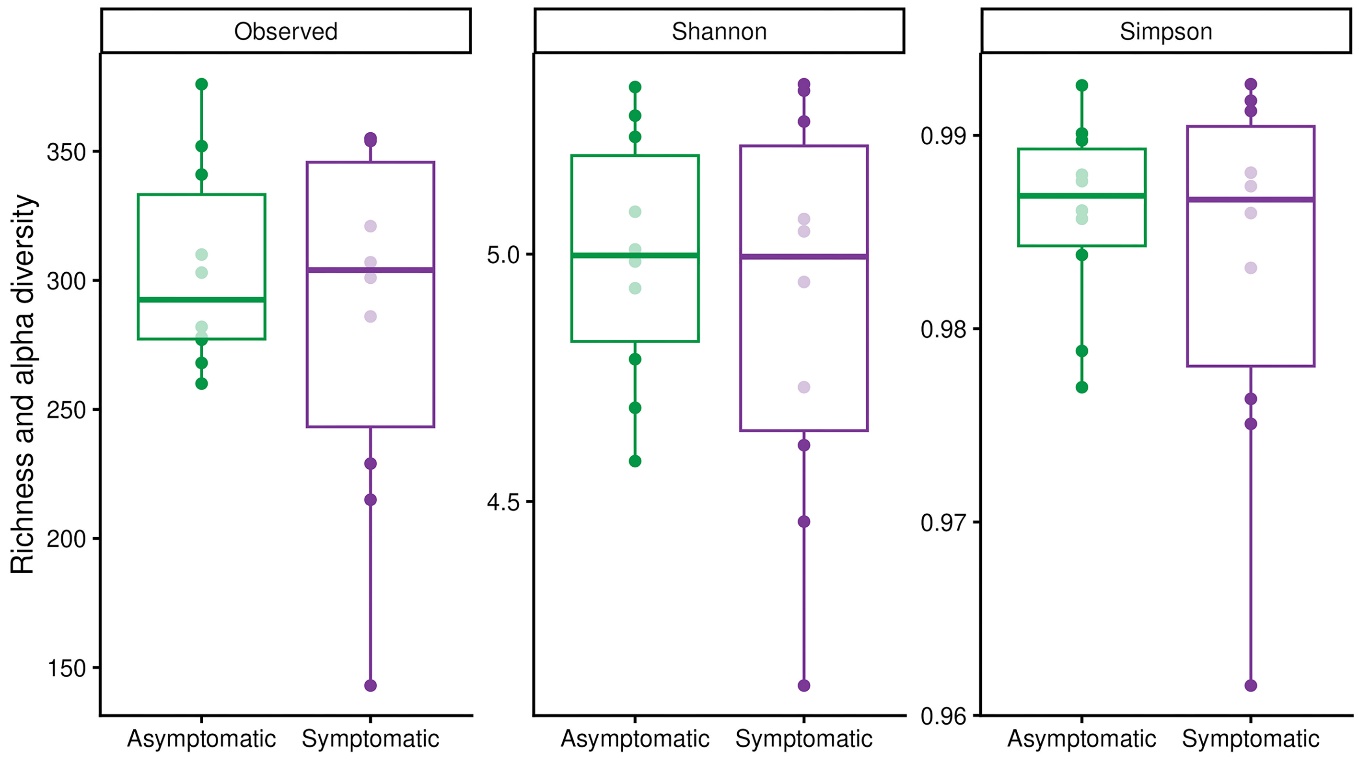
